# Overexposure to apoptosis via disrupted glial specification perturbs *Drosophila* macrophage function and reveals roles of the CNS during injury

**DOI:** 10.1101/2020.03.04.977546

**Authors:** Emma Louise Armitage, Hannah Grace Roddie, Iwan Robert Evans

## Abstract

Apoptotic cell clearance by phagocytes is a fundamental process during development, homeostasis and the resolution of inflammation. However, the demands placed on phagocytic cells such as macrophages by this process, and the limitations these interactions impose on subsequent cellular behaviours are not yet clear. Here we seek to understand how apoptotic cells affect macrophage function in the context of a genetically-tractable *Drosophila* model in which macrophages encounter excessive amounts of apoptotic cells. We show that loss of the glial transcription factor *repo*, and corresponding removal of the contribution these cells make to apoptotic cell clearance, causes macrophages in the developing embryo to be challenged with large numbers of apoptotic cells. As a consequence, macrophages become highly vacuolated with cleared apoptotic cells and their developmental dispersal and migration is perturbed. We also show that the requirement to deal with excess apoptosis caused by a loss of *repo* function leads to impaired inflammatory responses to injury. However, in contrast to migratory phenotypes, defects in wound responses cannot be rescued by preventing apoptosis from occurring within a *repo* mutant background. In investigating the underlying cause of these impaired inflammatory responses, we demonstrate that wound-induced calcium waves propagate into surrounding tissues, including neurons and glia of the ventral nerve cord, which exhibit striking calcium waves on wounding, revealing a previously unanticipated contribution of these cells during responses to injury. Taken together these results demonstrate important insights into macrophage biology and how *repo* mutants can be used to study macrophage-apoptotic cell interactions in the fly embryo.

Furthermore, this work shows how these multipurpose cells can be ‘overtasked’ to the detriment of their other functions, alongside providing new insights into which cells govern macrophage responses to injury in vivo.

## Introduction

Understanding the interactions between macrophages and apoptotic cells is an important biological question: failures in how immune cells deal with apoptotic cell death can lead to damaging autoimmune conditions [1]. Apoptotic cell clearance also plays a critical role in the resolution of inflammation and reprogramming immune cells during this process [2]. Furthermore, interactions between dying cells and macrophages occur at numerous sites of pathology including at sites of atherosclerosis [3] and in the chronically inflamed lungs of COPD patients [4]. As such, macrophage-apoptotic cell interactions have the potential to impact these diseases and many other damaging human conditions [1].

*Drosophila* has proven an excellent organism with which to study innate immunity [5], haematopoiesis [6] and blood cell function [7]. *Drosophila* blood is dominated by macrophage-like cells (plasmatocytes), with these cells making up 95% of the blood cells (hemocytes) in the developing embryo [8]. Embryonic macrophages disperse over the entire embryo during development, phagocytosing apoptotic cells and secreting matrix as they migrate [7]. Alongside their functional and morphological similarities to vertebrate macrophages, *Drosophila* macrophages are specified through the action of related transcription factors to those used in vertebrate haematopoiesis [9]. Failed dispersal or ablation of these embryonic macrophages leads to developmental abnormalities and failure to hatch to larval stages [10–12]. *Drosophila* macrophages are also able to respond to injury, mounting inflammatory responses to epithelial wounds [13]. In embryos and pupae, wounding elicits a rapid calcium wave through the epithelium, a process that requires transient receptor potential (Trp) channel function [14,15]. The increase in cytoplasmic calcium drives activation of Dual Oxidase (DUOX) and production of hydrogen peroxide, which is necessary for immune cell recruitment [14], resembling events upon tissue damage in higher organisms such as zebrafish [16,17].

In addition to macrophages, the other main phagocyte population within *Drosophila* embryos is the glia of the developing nervous system. *Drosophila* embryonic macrophages interact with the developing ventral nerve cord (VNC) and the glial cells that encase it, as they disperse along the ventral side of the embryo [18]. The VNC contains two populations of glia, the Sim-positive midline glia that help establish the ladder-like structure of neurons early in development and Repo-positive lateral glia [19]. Repo is a homeodomain transcription factor that specifies glial fate [20–22] and is required for expression of at least some phagocytic receptors in these cells [23]. While there are transcription factors in common between blood cells and glia (e.g. Gcm and Gcm2) [24,25], they are derived from distinct progenitors and Repo antagonises haematopoiesis to promote a glial fate [26]. Nonetheless both phagocytic populations express a similar repertoire of receptors for apoptotic cells, including Draper and Simu [27]. The close interactions between these glia and macrophages means that, should one population fail to clear apoptotic cells, it is likely that the other population would be able to both detect this deficiency and be able to compensate. Loss of apoptotic cell receptors such as Simu leads to a build up of apoptotic cells within developing embryos [28] and this impairs macrophage behaviours, including their developmental dispersal and inflammatory responses [29]. However, whilst mutants such as *simu* enable exposure of macrophages to elevated numbers of apoptotic cells in vivo, this approach is not ideal, as it also perturbs receptors that may be involved in immune cell reprogramming [30,31].

To investigate interactions between macrophages and apoptotic cells in more detail, we stimulated macrophages with enhanced levels of apoptotic cells using a genetic approach that did not alter the macrophages themselves. By impairing glial differentiation using *repo* mutants, we increased the number of apoptotic cells macrophages face within the developing embryo. This enhanced apoptotic challenge impaired macrophage dispersal, migration and their inflammatory responses to wounds. In this background, clearance of apoptotic cells by macrophages is not perturbed and migration can be rescued by preventing apoptosis. Surprisingly, and in contrast to phagocytic receptor mutants, blocking apoptotic cell death in the presence of defective glia failed to rescue wound responses. Further analysis revealed that injury-induced calcium waves propagate beyond the wounded epithelium and that this process is defective in *repo* mutants. This suggests that glial cells play an active role in the propagation of even the earliest responses to wounding. Thus, this model provides a unique insight into how macrophage-apoptotic cell interactions dictate macrophage responses to injury and the cell types that contribute to the activation of those responses.

## Materials and methods

### Fly lines and husbandry

Flies were reared at 25°C on cornmeal/agar/molasses media (see Supplementary Table 1 for recipe). *Srp-GAL4* [32] and *crq-GAL4* [13] were used to label *Drosophila* macrophages in the embryo in combination with *UAS-GFP* and/or *UAS-red stinger* [33]; *e22c-GAL4* (VNC and epithelium) [34], *act5C-GAL4* (ubiquitous) [35], *da-GAL4* (ubiquitous) [36], *elav-GAL4* (neuronal) [37] and *repo-GAL4* (glial cells) [38] were used to drive expression in other tissues. *UAS-GCaMP6M* [39] was used to image changes in cytoplasmic calcium concentration. Experiments were conducted on a *w^1118^* background and the following mutant alleles were used: *Df(3L)H99* [40], *repo^03702^* [20–22], *simu^2^* [28]. See Supplementary Table 2 for a full list of genotypes used in this study and sources of the *Drosophila* lines used. Embryos were collected from apple juice agar plates on which flies had laid overnight at 22°C. Embryos were washed off plates with distilled water and dechorionated in bleach for 1-2 minutes. Bleach was thoroughly washed away with distilled water ahead of fixation or mounting of embryos for live imaging. The absence of the fluorescent balancers *CTG*, *CyO dfd*, *TTG* and *TM6b dfd* [41,42] was used to select homozygous mutant embryos after dechorionation.

### Fixation and immunostaining

Dechorionated embryos were fixed and stained as per Roddie et al., 2019 [29]. Antibodies were diluted in PATx (0.1% Triton-X100 (Sigma-Aldrich), 1% BSA (Sigma-Aldrich) in PBS (Oxoid, Thermo Fisher, MA, USA)). Rabbit anti-GFP (ab290 1:1000; Abcam, Cambridge, UK) or mouse anti-GFP (ab1218 1:200; Abcam) were used to detect GFP-labelled macrophages. Rabbit anti-cDCP-1 (9578S 1:1000; Cell Signaling Technologies), mouse anti-Repo (concentrate of clone 8D12 used at 1:1000; Developmental Studies Hybridoma Bank, University of Iowa, USA) or mouse anti-Futch (supernatant of clone 22C10 used at 1:200; Developmental Studies Hybridoma Bank) were also used as a primary antibodies. Goat anti-mouse or goat anti-rabbit secondary antibodies conjugated to AlexaFluor568, AlexaFluor488 (Invitrogen, Thermo Fisher) or FITC (Jackson Immunoresearch, Cambridge, UK) were used to detect primary antibodies; these were diluted from stock solutions made according to the recommendations of the supplier (1:400 in PATx). Stained embryos were stored in DABCO mountant (Sigma-Aldrich) and mounted on slides for imaging.

### Imaging and analysis

Immunostained embryos were imaged using a 40X objective lens (CFI Super Plan Fluor ELWD 40x, NA 0.6) on a Nikon A1 confocal. Embryos containing GFP-labelled macrophages were stained for GFP and cDCP-1 and the phagocytic index (number of cDCP-1 punctae per macrophage per embryo), a measure of apoptotic cell clearance, was calculated from at least ten macrophages on the ventral midline in each embryo as per Roddie et al., 2019 [29].

Developmental dispersal was assessed by counting numbers of segments lacking GFP-labelled macrophages on the ventral side of the VNC in stage 13/14 embryos. Embryos that had been fixed and stained for GFP were scored using a Leica MZ205 FA fluorescent dissection microscope with a PLANAPO 2X objective lens. The same embryos were orientated ventral-side-up, imaged on a Nikon A1 confocal, and numbers of macrophages between epithelium and VNC counted (segments 4-8 were scored).

For live imaging of macrophage morphology, migration speed and wound responses and calcium dynamics upon injury, live embryos were mounted in voltalef oil (VWR) as per Evans et al., 2010 [18] and imaged on a Perkin Elmer UltraView Spinning Disk system using a 40X objective lens (UplanSApo 40x oil, NA 1.3). Epithelial wounds were made on the ventral side of embryos using a nitrogen-pumped Micropoint ablation laser (Andor, Belfast, UK), as per Evans et al., 2015 [43]. Macrophage inflammatory responses to wounds were imaged and analysed as per Roddie et al., 2019 [29]. Macrophage wound responses (numbers of macrophage at or touching the wound edge at 60 minutes divided by wound area in μm^2^) were normalised to control levels. The percentage of macrophages responding (% responders) was calculated by counting the proportion of macrophages present immediately following wounding that reached the wound within 60 minutes; macrophages already at the wound or absent from the field of view immediately after wounding were not considered in this analysis. Numbers of macrophages in the image taken prior to wounding (pre-wound) were counted as a measure of macrophages available to respond to the injury. The percentage of macrophages leaving the wound (% leavers) is the proportion of macrophages present at the wound site at any point during a 60-minute movie that leave the wound site; macrophages retaining contacts with the wound site or edge were not scored as leavers.

For analysis of random migration, 60-minute movies of GFP and red stinger double-labelled macrophages were made with macrophage position imaged every 2 minutes on the ventral midline between the overlying epithelium and VNC. Image stacks were despeckled and the movements of macrophages within maximum projections tracked as per Roddie et al., 2019 [29]. The manual tracking and Ibidi chemotaxis plugins were used to calculate speed per macrophage, per embryo, in Fiji [44].

Calcium dynamics were imaged using GCaMP6M expressed using a range of GAL4 lines. Calcium responses were quantified from average projections of z-stacks collected immediately before and after wounding. Initial wound responses of projections corresponding to epithelial z-slices (from *da-GAL4,UAS-GCaMP6M* embryos) were quantified by measuring the area of the GCaMP6M response and the corresponding mean gray value (MGV) of this region of interest (ROI) in Fiji immediately after wounding (F1). The same ROI was then used to measure the MGV of GCaMP6M fluorescence in the pre-wound image (F0). Both the area of the initial response and the F1/F0 ratio of MGVs were used as measures of the calcium response to injury (Figure 7c-d). For analysis of glial responses and the cells within the VNC more broadly (neurons and glia), *repo-GAL4* and *e22c-GAL4* were used to drive GCaMP6M expression, respectively. Responding glial cells were manually selected using the freehand selection tool in Fiji (Figure 8h-i), while the outer edge of the vitelline membrane was used to define a ROI to quantify GCaMP6M intensity for projections constructed from deeper volumes of the z-stack (Figure 8c).

### Image processing and statistical analyses

Images were despeckled in Fiji before maximum or average projections were made. All images were blinded ahead of quantification and Adobe Photoshop was used to assemble figures. Statistical analyses were performed in GraphPad Prism; legends and text contain details of statistical tests used. N numbers report numbers of individual *Drosophila* embryos analysed. Immunostained embryos are representative examples taken from batches of pooled embryos collected across multiple days.

## Results

### Embryonic macrophages disperse in close contact with the developing central nervous system

*Drosophila* embryonic macrophages migrate out from the presumptive head region to disperse over the entire embryo during development [7]. Migration along the developing ventral nerve cord (VNC) is an essential route for macrophage dispersal, with macrophages contacting the overlying epithelium and glial cells on the surface of the nerve cord (Figure 1a-b). During dispersal, macrophages encounter and clear apoptotic cells (Figure 1c), while VNC glia also phagocytose dying cells [45]. The interaction of macrophages and glia suggested to us that impairing glial-mediated apoptotic cell clearance could increase exposure of macrophages to apoptotic cell death in vivo, thus providing a model with which apoptotic cell-macrophage interactions and their effects on macrophage behaviour could be studied in detail.

**Figure 1.**
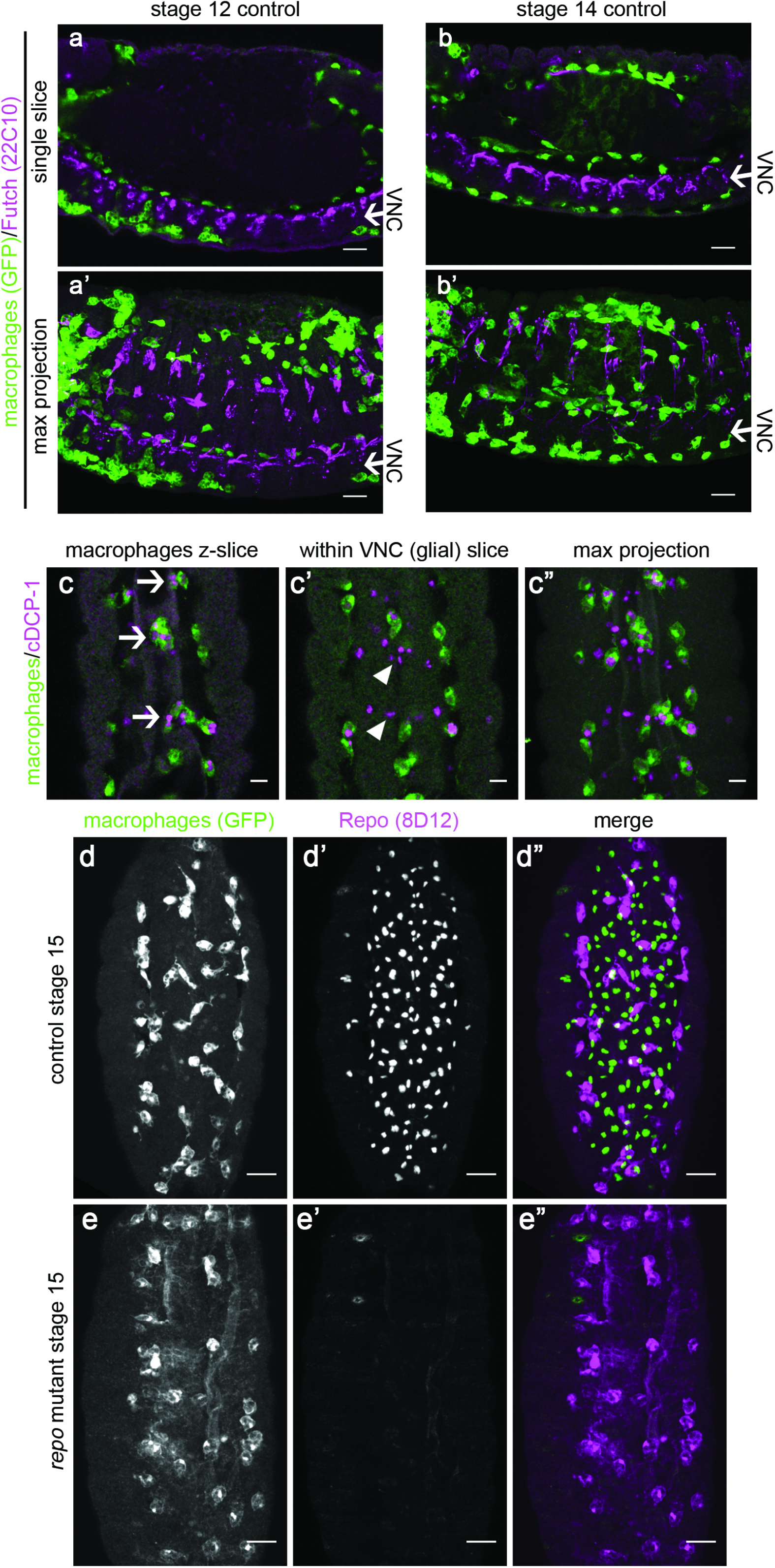
Interaction of macrophages and glial cells during *Drosophila* embryonic development. (a-b) single z-slices (a,b) and maximum projections (a’,b’) of immunostained control embryos with GFP-labelled macrophages (green) showing progression of macrophages along both sides of the ventral nerve cord (VNC, Futch staining; purple) at stage 12 (a) and 14 (b). Arrows indicate position of VNC; embryos are laterally orientated with anterior to the left, and ventral down. (c) ventral views of a stage 15 control embryo immunostained for apoptotic cells (anti-cDCP-1, purple) and macrophages (anti-GFP, green); panels show single z-slices showing engulfment of apoptotic cells by macrophages (arrows, c) and apoptotic cells within the VNC inaccessible to macrophages (arrowheads, c’), and a maximum projection of this region (c”). (d-e) ventral views of stage 15 control and *repo* mutant embryos immunostained for GFP to show macrophages (d,e) and anti-Repo (d’,e’); macrophages and Repo are green and purple, respectively, in merged images (d”, e”). All scale bars represent 10μm. Genotypes are as follows: *w;srp-GAL4,UAS-GFP/+;crq-GAL4,UAS-GFP/+* (a-b), *w;;crq-GAL4,UAS-GFP* (c-d), *w;;repo^03702^,crq-GAL4,UAS-GFP* (e).

### Increased efferocytosis by embryonic macrophages in the absence of functional glial cells

Both macrophages and glial cells use phagocytic receptors such as Draper and Simu to clear apoptotic cells [28,46,47]. The absence of these receptors perturbs clearance of apoptotic cells, which in turn disrupts macrophage functions, including migration speed and inflammatory responses [29,43]. While this suggests that apoptotic cells modulate macrophage behaviour in flies, these interventions remove genes with key roles in pathways that control macrophage fate [30]. Therefore, in order to expose macrophages to increased amounts of apoptotic cell death without directly impacting their own specification mechanisms or removing important regulators of phagocytosis, we targeted the glia of the VNC. We hypothesised that blocking glial-mediated clearance would lead to macrophages becoming exposed to increased numbers of apoptotic cells due to the inability of glial cells to contribute to this process.

The homeodomain transcription factor Repo is expressed by all glial cells within the developing VNC [20–22], except the midline glia, which are specified via the action of *sim* [48]. Repo is absolutely required for normal specification of these glial cells and absence of *repo* leads to the failure to express a variety of phagocytic receptors required for apoptotic cell clearance [23]. Repo is not expressed by macrophages in the developing embryo (Figure 1d; [38]) and *repo^03702^* mutants lack detectable protein expression in the VNC (Figure 1e).

To test whether failed glial specification would lead to macrophages encountering increased numbers of apoptotic cells in the developing embryo, we analysed macrophage morphology. Macrophages in *repo* mutants are highly vacuolated compared to controls (Figure 2a-b); vacuoles within *Drosophila* embryonic macrophages typically contain previously engulfed apoptotic cells [49]. To test whether apoptotic cell clearance by macrophages is increased in *repo* mutants, control and *repo* mutant embryos were immunostained for a marker of apoptotic cell death (cleaved DCP-1 immunostaining; DCP-1 is cleaved by caspases during apoptosis [50]). Macrophages in *repo* mutants contain far higher numbers of cDCP-1 positive inclusions compared to controls (Figure 2c-e). Furthermore, macrophages can be labelled using *crq-GAL4* or *srp-GAL4* in a *repo* mutant background and efficiently engulf apoptotic cells (Figure 2), behaviour consistent with their normal specification and physiology. These results therefore suggest that loss of *repo* function is a suitable tool via which the effects of increased macrophage-apoptotic cell contact can be analysed in vivo.

**Figure 2.**
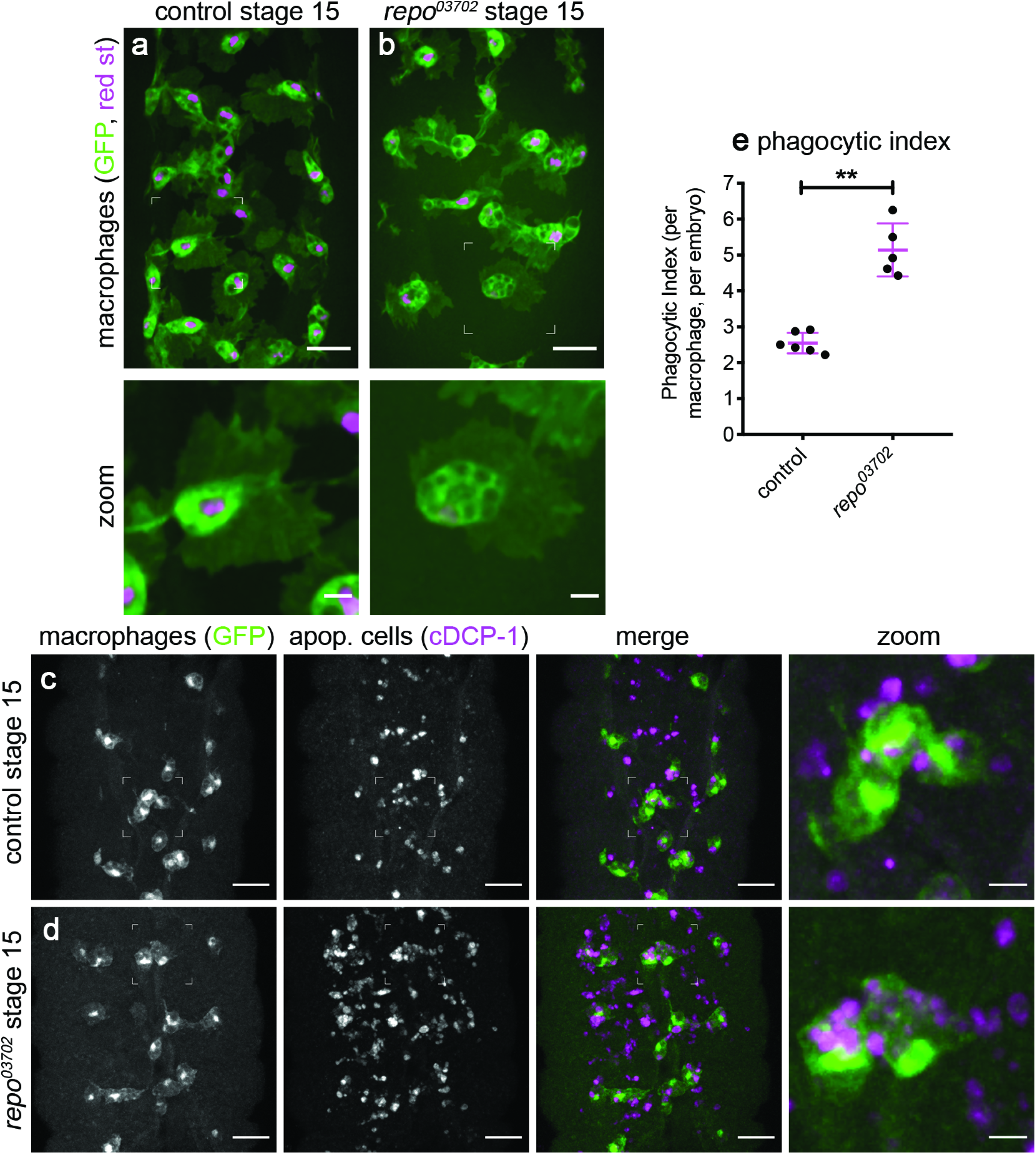
Increased macrophage-mediated apoptotic cell clearance in the absence of glial specification. (a-b) ventral views of stage 15 control (a) and *repo* mutant (b) embryos containing GFP and red stinger-expressing macrophages; lower panels show zooms of macrophages indicated by boxes in upper panels. (c-d) ventral views of stage 15 control (c) and *repo* mutant (d) embryos containing GFP-labelled macrophages immunostained for GFP (green in merge) and apoptotic cells (anti-cDCP-1, purple in merge); zooms show macrophages indicated in boxed regions. (e) scatterplot of phagocytic index (cDCP-1 puncta per macrophage, per embryo); error bars and line show standard deviation and mean, respectively; p=0.0043 via Mann-Whitney test (n=6 and 5 for controls and *repo* mutants, respectively). Scale bars represent 20μm or 5μm in zooms; ** indicates p<0.001; genotypes are *w;srp-GAL4,UAS-GFP/srp-GAL4,UAS-red stinger* (a), *w;srp-GAL4,UAS-GFP/srp-GAL4,UAS-red stinger;repo^03702^* (b), *w;;crq-GAL4,UAS-GFP* (c, e) and *w;;repo^03702^,crq-GAL4,UAS-GFP* (d-e).

### An increased burden of apoptotic cell clearance is associated with impaired developmental dispersal of macrophages

Apoptotic cells represent the top priority for *Drosophila* macrophages to respond to within developing embryos [51]. As a result, increased numbers of apoptotic cells have the potential to disrupt macrophage behaviours, such as their dispersal and recruitment to sites of tissue injury. *repo* mutants lack gross dispersal defects, with macrophages present along both sides of the VNC at stage 13 (Figure 3a-b). However, reduced numbers of macrophages are present on the ventral midline in *repo* mutants (Figure 3c-e), suggesting an impairment in dispersal. Further increasing apoptotic burden in a *repo* mutant background (via removal of the apoptotic cell clearance receptor Simu) causes large dispersal defects (Figure 3a-b), comparable to those observed when critical regulators of migration are absent, e.g. SCAR/WAVE [49]. Taken together, these results indicate that challenge of macrophages with excessive amounts of apoptosis can impair developmental dispersal of macrophages.

**Figure 3.**
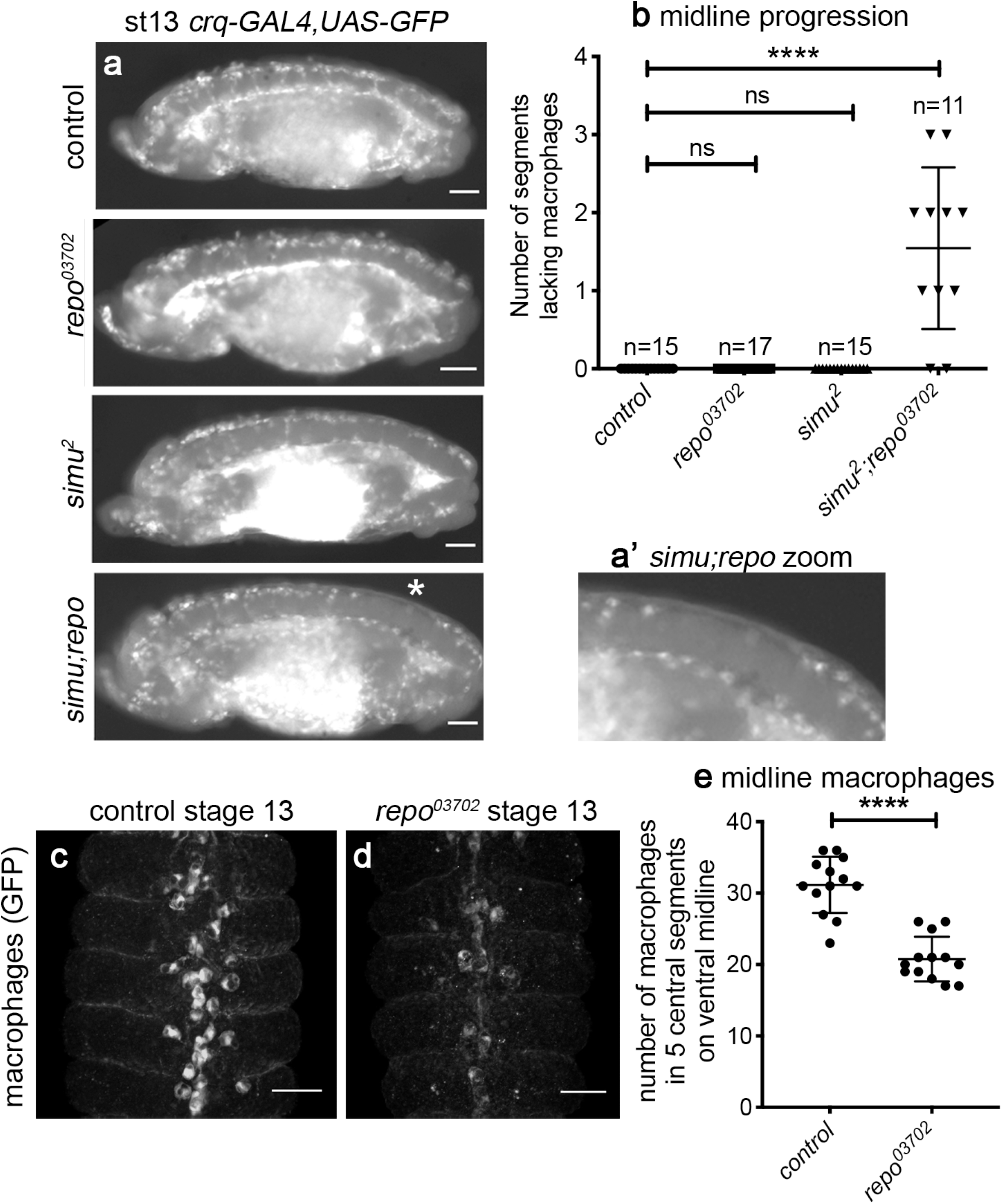
Excessive amounts of apoptotic cell death impair macrophage dispersal. (a) lateral views of stage 13/14 control, *repo*, *simu* and *simu;repo* double mutant embryos containing GFP-labelled macrophages. A complete line of macrophages is present on midline on the ventral side of the VNC in all genotypes, with the exception of *simu;repo* double mutants (gap indicated via an asterisk); (a’) shows zoom of this region. (b) scatterplot showing quantification of midline progression defects (numbers of segments lacking macrophages on the ventral side of the VNC); p<0.0001 (for *simu;repo* vs each other condition; no other comparisons are significantly different) via one-way ANOVA with a Tukey’s multiple comparison post-test; n=15 (controls), 17 (*repo* mutants), 15 (*simu* mutants) and 11 (*simu;repo*). (c-d) ventral views of stage 13 control and *repo* mutant embryos containing GFP-expressing macrophages (immunostained via anti-GFP). (e) scatterplot showing quantification of numbers of macrophages in five central segments on the ventral midline; p<0.0001 via Mann-Whitney test, n=13 (controls), 13 (*repo* mutants). Scale bars represent 50μm (a) and 25μm (c-d); lines and error bars on scatterplots show mean and standard deviation, respectively; **** indicates p<0.0001. Genotypes are *w^1118^;;crq-GAL4,UAS-GFP* (control), *w^1118^;;P{PZ}repo^03702^,crq-GAL4,UAS-GFP* (*repo^03702^*), *w^1118^;simu^2^;crq-GAL4,UAS-GFP* (*simu^2^*) and *w^1118^;simu^2^;P{PZ}repo^03702^,crq-GAL4,UAS-GFP* (*simu;repo*).

### Apoptotic cells are responsible for attenuation of macrophage migration in repo mutants

To test the effects of increased apoptotic cell exposure on macrophage migration, we tracked the movements of macrophages on the ventral midline at stage 15 after completion of developmental dispersal, a point at which macrophages exhibit wandering/“random migration”. In *repo* mutant embryos, macrophages move at significantly slower speeds compared to those in controls (Figure 4). In order to test whether this attenuation of migration speed is dependent on interactions with apoptotic cells, we removed all developmental apoptosis from a *repo* mutant background using the *Df(3L)H99* deficiency, which deletes the pro-apoptotic genes *Hid*, *reaper* and *grim* [40]. In these *repo* mutants that lack apoptosis there is a significant rescue of macrophage migration speed (Figure 4c-e), suggesting that it is interactions with apoptotic cells that impair macrophage migration in vivo, rather than defective glial specification per se.

**Figure 4.**
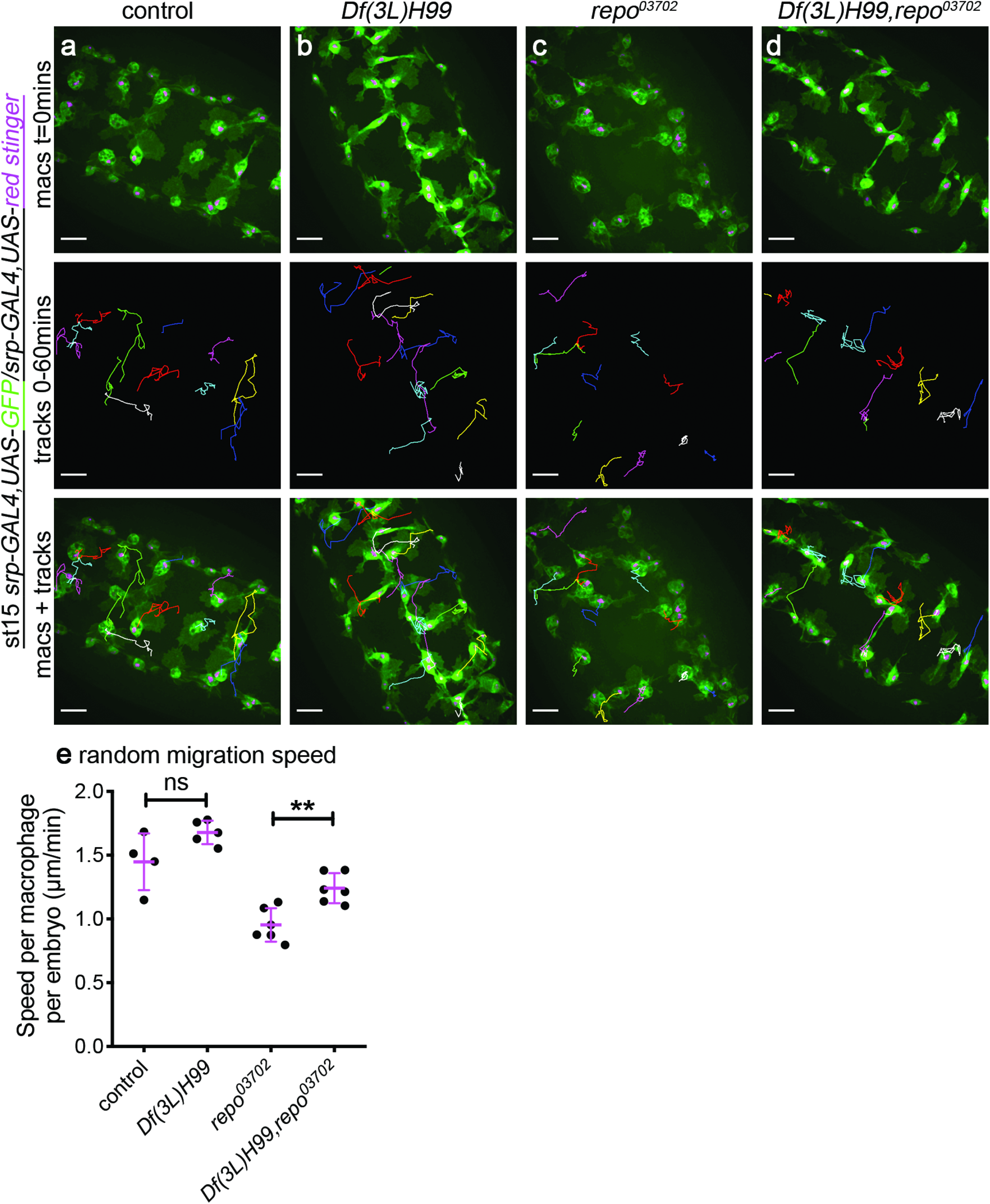
Excessive amounts of apoptotic cells impair macrophage migration. (a-d) ventral images of GFP and red stinger-labelled macrophages (green and purple, respectively) in stage 15 control (a), *Df(3L)H99* mutant (b), *repo^03702^* mutant (c) and *Df(3L)H99,repo^03702^* double mutant embryos (d). Upper panels show initial position of macrophages on the ventral midline; central panels show tracks taken from subsequent 60-minute period from the timepoint shown in upper panel; lower panel shows overlay of tracks over initial positions of macrophages. (e) scatterplot showing speed per macrophage per embryo (μm per min); lines and error bars show mean and standard deviation, respectively. Removing apoptosis from a *repo* mutant background (*Df(3L)H99;repo*) rescues migration speed (p=0.0043 via Mann-Whitne test); n= 4 (control), 5 (*Df(3L)H99*), 6 (*repo*) and 6 (*Df(3L)H99,repo*). Scale bars represent 20μm; ns and ** indicate not significant and p<0.01. Genotypes are *w^1118^;srp-GAL4,UAS-red stinger/srp-GAL4,UAS-GFP* (control), *w^1118^;srp-GAL4,UAS-red stinger/srp-GAL4,UAS-GFP;Df(3L)H99* (*Df(3L)H99*), *w^1118^;srp-GAL4,UAS-red stinger/srp-GAL4,UAS-GFP;P{PZ}repo^03702^* (*repo^03702^*) and *w^1118^;srp-GAL4,UAS-red stinger/srp-GAL4,UAS-GFP;Df(3L)H99,P{PZ}repo^03702^* (*Df(3L)H99,repo^03702^*).

### Functional glia are required for normal macrophage migration to wounds

Given the impaired migration of macrophages in *repo* mutant embryos, we analysed their inflammatory responses to sterile, laser-induced wounds. Wounding of *repo* mutant embryos revealed that reduced numbers of macrophages reached wound sites by 60-minutes post-wounding compared to controls (Figure 5a-b,d). Since there are fewer macrophages present locally at this developmental stage in *repo* mutants (Figure 5c), the percentage of cells responding to the injury was also quantified, revealing a significant decrease in the ability of cells to respond to wounds (Figure 5e; Supplementary Movie 1). Early migration away from wounds does not appear to underlie these defects (Figure 5f), suggesting that excessive amounts of apoptotic cells turn off wound responses in *repo* mutants. To further evidence this conclusion, we removed developmental apoptosis, comparing wound responses in *Df(3L)H99* mutants and *Df(3L)H99,repo* double mutant embryos. In contrast to the rescue of migration speed, removing apoptosis from a *repo* mutant background failed to rescue either the numbers of cells responding to the injury or the percentage of cells able to respond (Figure 6). Therefore, the underlying defect in *repo* mutants that hinders macrophage inflammatory responses does not appear to be contact with apoptotic cells (Figure 6), in contrast to the effect seen for migration defects (Figure 4).

**Figure 5.**
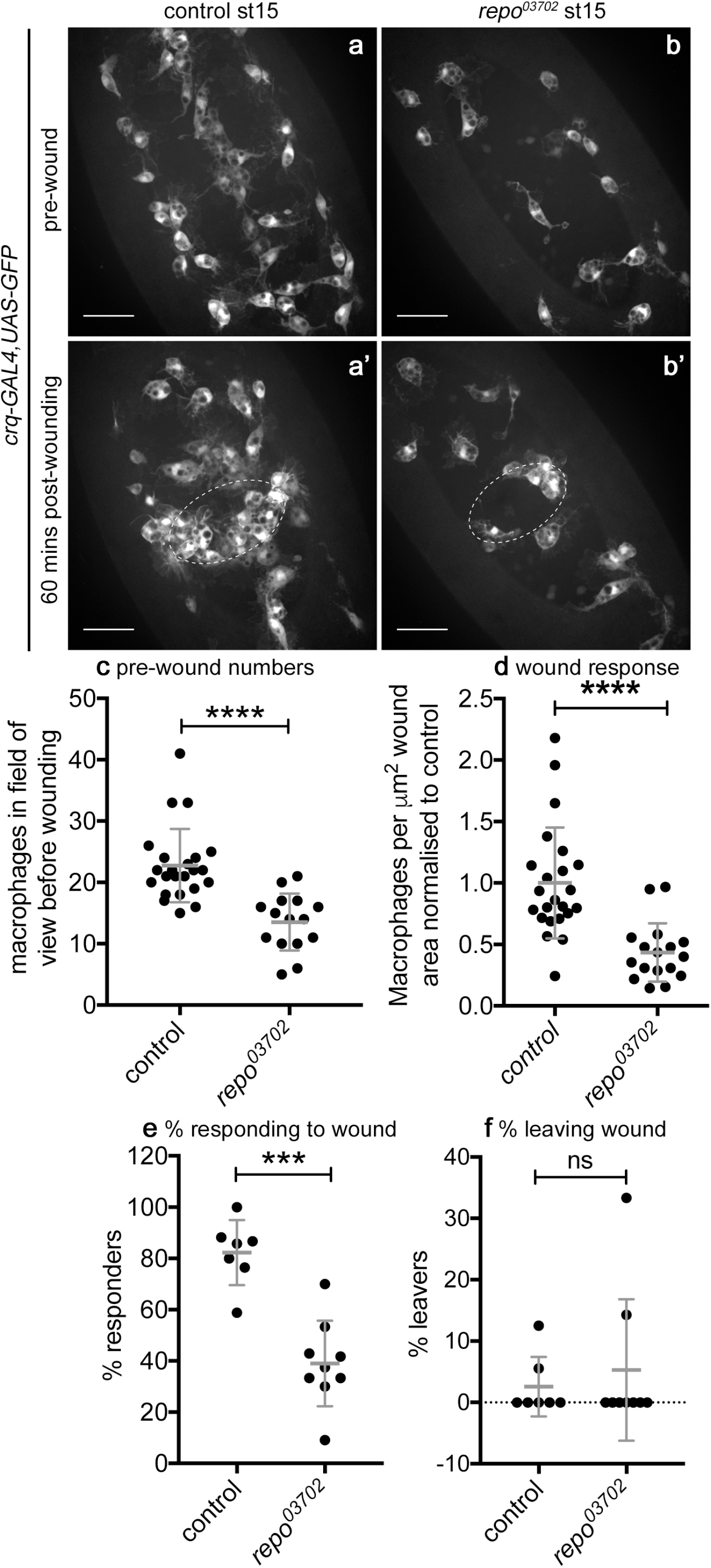
*repo* is required for normal macrophage inflammatory responses to wounding. (a-b) ventral views showing localisation of GFP-labelled macrophages on the ventral midline in control (a, a’) and *repo* mutant (b, b’) embryos immediately before wounding (a,b) and at 60-minutes post-wounding (a’,b’); white dotted ellipses indicate wound edges. (c) scatterplot showing numbers of macrophage in the field of view ahead of wounding per embryo; p<0.0001 via Mann-Whitney test (n=23 and 15 control and *repo*, respectively). (d) scatterplot showing wound responses quantified via density of macrophages at wounds, normalised to control; p<0.0001 via Mann-Whitney test (n=23 and 17 control and *repo*, respectively). (e) scatterplot showing percentage of macrophages responding to wounds; p=0.0003 via Mann-Whitney test (n=7 and 9 control and *repo*, respectively). (f) scatterplot showing percentage of macrophages that leave the wound (having initially migrated to the wound); p>0.999 via Mann-Whitney test (n=7 and 9 control and *repo*, respectively). Lines and error bars in scatterplots show mean and standard deviation, respectively; scale bars represent 20μm; ns, *** and **** indicate not significant (p>0.05), p<0.001 and p<0.0001, respectively. Genotypes are *w;;crq-GAL4,UAS-GFP* (control) and *w;crq-GAL4,UAS-GFP,repo^03702^* (*repo^03702^*).

**Figure 6.**
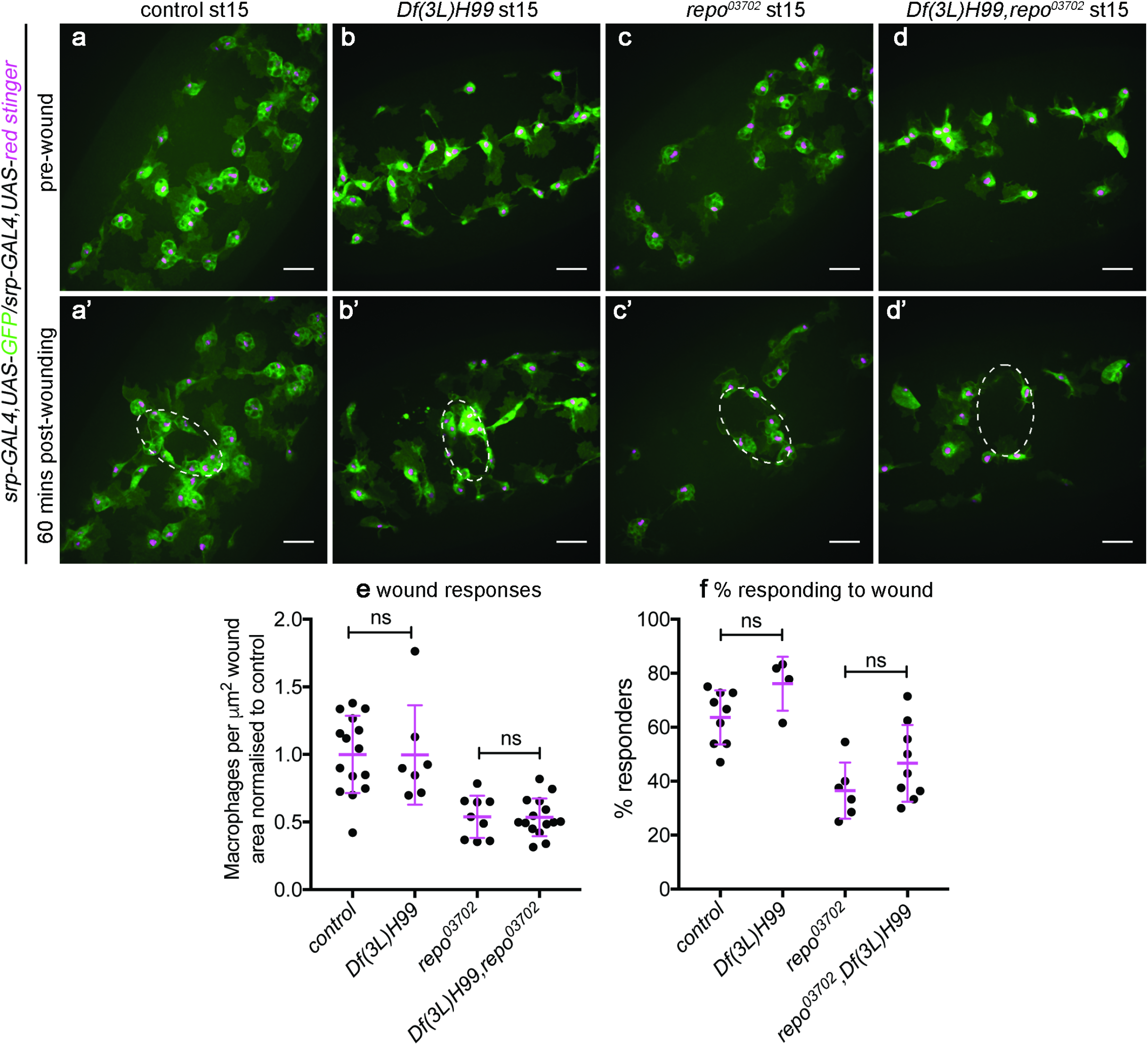
Loss of *repo* impairs wound responses but contact with apoptotic cells is not the overriding cause of this defect. (a-b) ventral views showing localisation of GFP and red stinger-labelled macrophages on the ventral midline in control (a,a’), *Df(3L)H99* (b,b’), *repo* (c,c’) and *Df(3L)H99,repo* double mutant embryos (d,d’) immediately before wounding (a-d) and at 60-minutes post-wounding (a’-d’); white dotted ellipses indicate wound edges. (e) scatterplot showing wound responses quantified via density of macrophages at wounds, normalised to control; Mann-Whitney tests used to compare control vs *Df(3L)H99* (n=15 and 7, respectively; p=0.63) and *repo* vs *Df(3L)H99,repo* (n=9 and 15, respectively; p>0.99). (f) scatterplot showing percentage of macrophages responding to wounds; Mann-Whitney tests used to compare control vs *Df(3L)H99* (n=9 and 4, respectively p=0.055) and *repo* vs *Df(3L)H99,repo* (n=6 and 6, respectively; p=0.17). Lines and error bars in scatterplots show mean and standard deviation, respectively; scale bars represent 20μm; ns indicates not significant (p>0.05). Genotypes are *w;srp-GAL4,UAS-GFP/srp-GAL4,UAS-red stinger* (control), *w;srp-GAL4,UAS-GFP/srp-GAL4,UAS-red stinger;Df(3L)H99* (*Df(3L(H99)*), *w;srp-GAL4,UAS-GFP/srp-GAL4,UAS-red stinger;repo^03702^* (*repo^03702^*) and *w;srp-GAL4,UAS-GFP/srp-GAL4,UAS-red stinger;Df(3L)H99,repo^03702^* (*Df(3L)H99,repo^03702^*).

### Neurons and glia respond to injury through changes in their calcium dynamics

Since removing the apoptotic cell burden from macrophages in *repo* mutants failed to improve their inflammatory responses to wounds (Figure 6), we investigated whether upstream signalling mechanisms that form part of the normal response to wounds remain intact in this mutant background. Laser ablation of embryos triggers the rapid spread of a calcium wave through the epithelium, away from the site of damage (Figure 7a; Supplementary Movie 2; [14]). This wave of calcium activates hydrogen peroxide production via DUOX, which is itself required for migration of macrophages to wound sites [14,43,52]. Imaging calcium dynamics after wounding using the cytoplasmic calcium sensor GCaMP6M [39], we were unable to identify a difference immediately post-wounding in the epithelial calcium responses of *repo* mutant embryos compared to controls (Figure 7a-d; Supplementary Movie 2). However, in performing these experiments we noticed that the calcium response was not limited to the epithelium, with this signal visible deeper within the embryo including within the neurons and glia of the VNC (Figure 7e-g) – a structure located immediately underneath the epithelium on the ventral side of the embryo (Figure 1a-b). These changes in intracellular calcium were not limited to the damaged tissue and extended away from the necrotic core of the wound (Figure 7e-g; Supplementary Movie 3), with changes in calcium levels particularly striking within the axons of the CNS (Figure 7g). Strikingly, quantification of calcium responses in sub-epithelial regions upon wounding showed a reduced calcium response in *repo* mutants compared to controls (Figure 8a-c), suggesting that glial cells are responsive to injury and contribute to damage-induced signalling.

**Figure 7.**
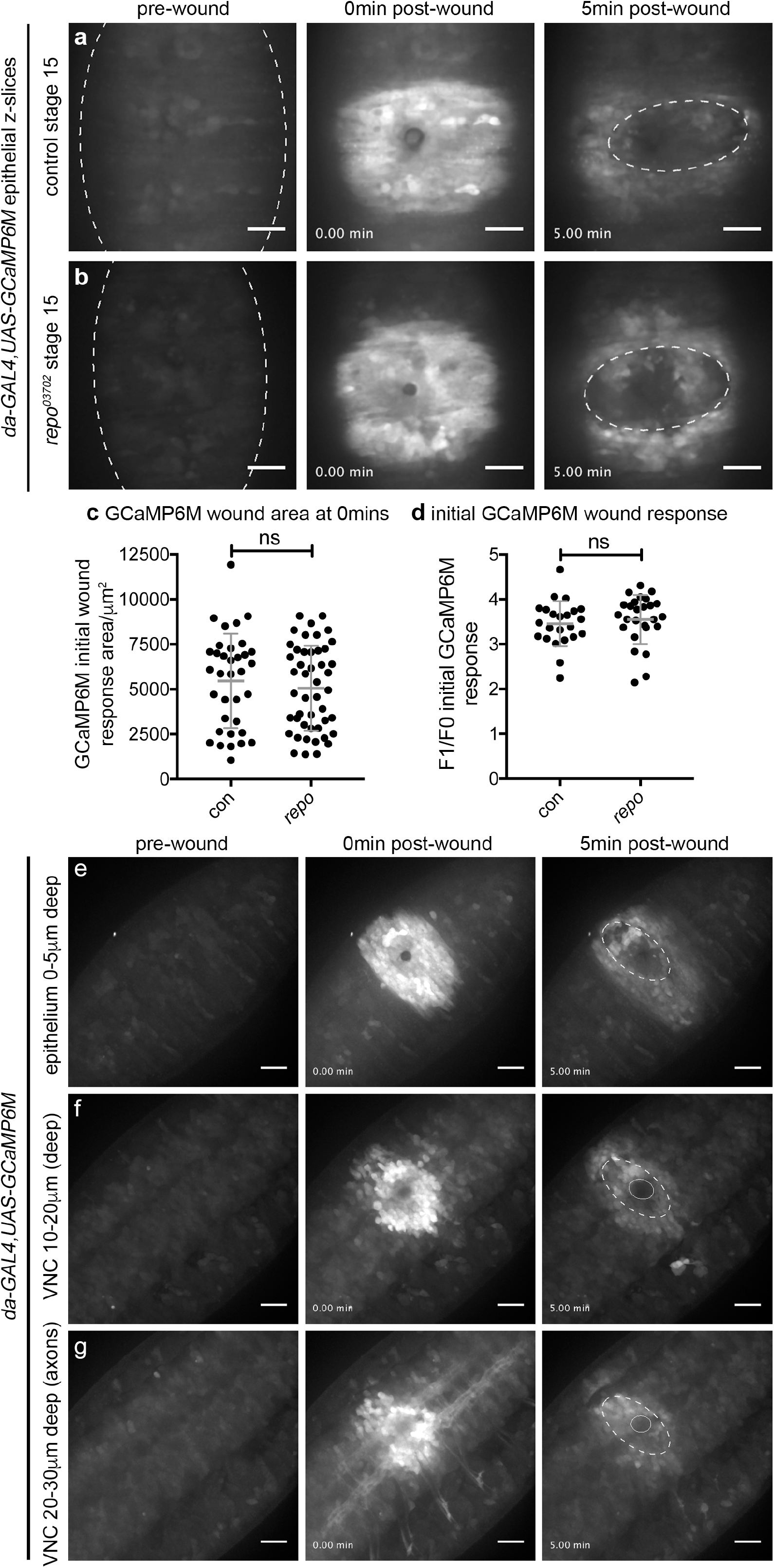
Defective calcium responses to wounding in *repo* mutants are restricted to less superficial tissues. (a-b) calcium levels imaged via ubiquitous expression of GCaMP6M (*da-GAL4,UAS-GCaMP6M*) on wounding of the ventral surface of control (a) and *repo* mutant (b) stage 15 embryos; images show pre-wound calcium levels, immediately after wounding (0min) and 5 minutes after wounding. Dotted lines show edges of embryo and wound edges. (c) scatterplot showing area of GCaMP6M response immediately after wounding; Mann-Whitney test used to compare control vs *repo* (n=35 and 47, respectively; p=0.66). (d) scatterplot showing ratio of GCaMP6M intensity in wound area before and immediately after wounding; Mann-Whitney test used to compare control vs *repo* (n=23 and 26, respectively; p=0.22). (e-g) maximum projections of superficial/epithelial regions (0-5μm, e), superficial half of the VNC (10-20μm from surface, f) and deeper/dorsal half of the VNC (20-30μm from the surface, g) of the ventral side of a wounded control embryo containing ubiquitous expression of GCaMP6M; panels show pre-wound calcium levels, immediately after wounding (0min) and 5 minutes after wounding. Dotted lines show edges of epithelial wound at 5 minutes, solid circles show physical damage at deeper regions of the embryo; embryo has not been orientated in order to show larger region of the response to wounding. Lines and error bars in scatterplots show mean and standard deviation, respectively; scale bars represent 20μm; ns indicates not significant (p>0.05). Genotypes are *w;;da-GAL4,UAS-GCaMP6M* (control) and *w;;repo^03702^,da-GAL4,UAS-GCaMP6M* (*repo*).

### *repo* is required for normal calcium responses to injury in the developing embryo

Using the tissue-specific drivers *elav-GAL4* (neurons) and *repo-GAL4* (glia) to express GCaMP6M confirmed that both neurons and glial cells within the VNC respond to injury by transiently increasing their cytosolic calcium levels (Figure 8d-e; Supplementary Movies 4 and 5). Changes in cytoplasmic calcium are transmitted beyond the confines of the wound (Figure 8d,g) and remain elevated at wound sites several hours post-injury (Supplementary Figure 1). These responses suggest that these cells communicate damage signals away from sites of tissue damage, potentially contributing to inflammatory recruitment of macrophages and regenerative processes. Additionally, the transmission of calcium signals distal to the site of physical injury suggests these responses are not simply limited to those cells damaged during the wounding process.

In *repo* mutants, while there is less glial proliferation, residual glial cells are present, but lack late glial markers [20,21] and normal patterns of phagocytic receptor expression [23]. In contrast to the wild-type situation, wounding of *repo* mutants with glial-specific expression of GCaMP6M showed a reduction in the calcium response on injury (Figure 8e-f,h-i). Taken together, these data indicate that, in contrast to our present understanding, tissues other than the epithelium undergo alterations in calcium signalling upon injury. Furthermore, the reduced calcium responses observed in *repo* mutants on wounding may therefore contribute to the reduced inflammatory responses undertaken by macrophages.

**Figure 8.**
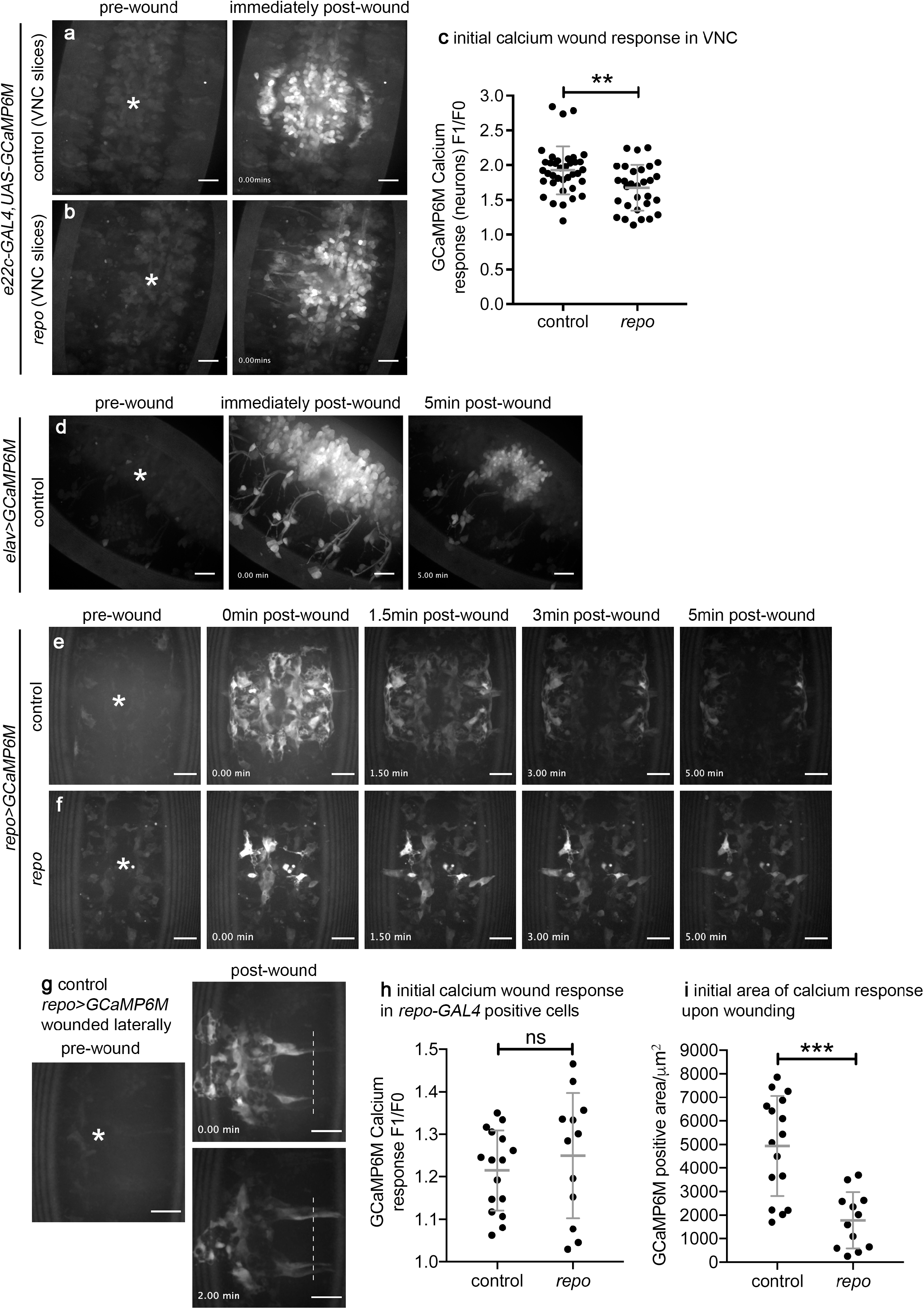
Non-epithelial tissues contribute to wound responses. (a-b) calcium levels within the VNC imaged via expression of *UAS-GCaMP6M* via *e22c-GAL4* (projections assembled from slice range 30-10μm deep from surface of the embryo) on wounding of the ventral surface of control (a) and *repo* mutant (b) stage 15 embryos; images show pre-wound calcium levels and immediately after wounding (0min). (c) scatterplot showing GCaMP6M response within VNC immediately after wounding (F1/F0) of control and *repo* mutants labelled using *e22c-GAL4,UAS-GCaMP6M*; Mann-Whitney test used to compare control vs *repo* (n=37 and 30, respectively; p=0.0051). (d) calcium levels in neuronal cells (labelled using *elav-GAL4* to drive *UAS-GCaMP6M* expression) on wounding of the ventral surface of a stage 15 embryo; images show pre-wound calcium levels, immediately after wounding (0min) and 5-minutes after wounding. (e-g) calcium levels in glial cells (labelled using *repo-GAL4* to drive *UAS-GCaMP6M* expression) on wounding of the ventral surface of control (e,g) and *repo* (f) mutant stage 15 embryos; images show pre-wound calcium levels, immediately after wounding (0min) and 5-minutes after wounding. (g) shows embryo wounded more laterally and subsequent spread of calcium signal along glial cells in more lateral positions; dotted line shows equivalent position in the 0 minute and 2 minute post-wound timepoint images. (h-i) scatterplots showing GCaMP6M responses in glial cells immediately after wounding (h, F1/F0) and the initial area (μm^2^) of the GCaMP6M response in control and *repo* mutant embryos labelled via *repo-GAL4,UAS-GCaMP6M* (i); Mann-Whitney test used to compare control vs *repo* (n=16 and 12, respectively; p=0.42 (h) and p=0.0003 (i)). Asterisks show position of wounds in pre-wound images; scale bars represent 20μm; lines and error bars show mean and standard deviation in all scatterplots; **, *** and ns denote p<0.01, p<0.001 and not significant (p>0.05), respectively. Genotypes are *w;e22c-GAL4,UAS-GCaMP6M* (a,c), *w;e22c-GAL4,UAS-GCaMP6M;repo^03702^* (b-c), *w,elav-GAL4/w;UAS-GCaMP6M/+* (d), *w;repo-GAL4,UAS-GCaMP6M* (e,g,h-i) and *w;repo-GAL4,UAS-GCaMP6M;repo^03702^* (f,h-i).

## Discussion

Clearance of apoptotic cells is associated with reprogramming of phagocytes such as macrophages towards anti-inflammatory states and is part of the process of resolution of inflammation [2]. Here we show that by impairing glial specification using *repo* mutants, macrophages can be challenged with increased levels of apoptotic cell death in vivo. Challenging ‘wild-type’ macrophages in this way causes them to become engorged with phagocytosed apoptotic cells. The excessive amounts of apoptosis are associated with impaired developmental dispersal, slowed migration speeds and attenuated inflammatory responses. While migration speeds can be rescued by removing apoptosis from a *repo* mutant background, wound responses are not improved, suggesting interactions with apoptotic cells do not represent the primary cause of this phenotype. Instead, in contrast to our current understanding, tissues in addition to the wounded epithelium are wound-responsive and glia and the cells they support are likely to contribute to immune cell recruitment and repair mechanisms in *Drosophila*.

Macrophages in *repo* mutants exhibit impaired dispersal and reduced migration speeds, phenotypes consistent with other *Drosophila* mutants that perturb apoptotic cell clearance, including *SCAR/WAVE* [49], *draper* [43] and *simu* [29]. Similarly, removal of the apoptotic clearance burden, via ablation of the apoptotic machinery, improves migration speeds and suggests that apoptotic cells are responsible for these phenotypes. Excessive production of find-me cues – chemoattractants released from apoptotic cells to alert phagocytes to their presence [53] – represent one potential explanation, with such cues distracting or overwhelming macrophages. Consistent with this, apoptotic cells have been inferred to be the most highly-prioritised migratory cue for macrophages in the fly embryo [51]. Currently, this remains hard to investigate further, since the nature and identity of find-me cues remains to be discovered in this organism. Alternatively, reprogramming of macrophages to a different transcriptional or activation state may also explain these phenotypes [54], although little evidence for macrophage polarisation comparable to that observed in vertebrate myeloid cells currently exists for *Drosophila*.

Phagocytes such as *Drosophila* macrophages may have to decide whether to engulf or move: for instance both dendritic cells and *Dictyostelium* amoebae pause during macropinocytosis [55,56], a process related to phagocytosis. Similarly, lysosomal storage disorders that lead to vacuolation are associated with perturbed immune cell migration in patient-derived cells [57–59] and experimental models [60]. Sequestration of regulators of both phagocytosis and motility, such as the lysosomal Trp channel Trpml (the fly homologue of TRPML1/MCOLN1) [61], by excessive phagocytic cup formation in the face of elevated apoptotic cell burdens, could antagonise the ability of phagocytes to migrate. *repo* mutants represent a model to further understand how apoptotic cells regulate changes in macrophage behaviour in vivo, such as their migration and phagocytic capacities, especially since initial specification of macrophages does not appear compromised in this background and *repo* plays no direct functional role in these cells.

Macrophage inflammatory responses were not improved by blocking apoptosis in *repo* mutants. This was surprising, since preventing apoptosis in *simu* mutants, another background in which macrophages face large numbers of apoptotic cells, did improve macrophage responses to injury [29]. Thus it is unlikely that apoptotic cell-macrophage interactions represent the primary cause of wound recruitment defects in *repo* mutants. Faster and more sensitive imaging technologies, alongside the use of alternative drivers to express genetically-encoded calcium reporters enabled capture of larger volumes of the embryo during wounding in comparison to previous studies [14]. This revealed that those tissues surrounding the damaged epithelium, such as the neurons and glia of the VNC, were also responsive to injury. Loss of *repo* function causes defects in glial specification, proliferation [20–22] and also leads to a perturbed calcium response in the VNC and glia. Potentially, *repo* drives a transcriptional program enabling glia to respond to injury. Failed glial specification may have a “knock-on” effect and hinder the damage responses of neurons, which are supported by glial cells. Alternatively, the decreased number or mispositioning of glial cells in *repo* mutants [20] may decrease the amplitude or spread of wound signals activated by injury, such as via defective cell-cell contacts between glia. Impairing the ability of surrounding tissues to respond to injury may therefore perturb the generation of wound cues required for normal macrophage migration to sites of tissue damage and could potentially impact evolutionarily-conserved CNS repair processes [62], especially since calcium waves in the developing *Drosophila* wing disc are required for regeneration following mechanical injury [63].

The use of calcium as a regulator of wound responses is conserved across evolution with calcium waves visible upon transection of zebrafish larval fins [17], and immediately after wounding *Xenopus* embryos [64] and the *C. elegans* epidermis [65]. Release of calcium from internal stores during wave propagation is also a commonality in these models. The mechanism of activation in the *Drosophila* CNS remains to be established, but in zebrafish tailfin wounds the release of ATP from damaged cells may activate P2Y receptors, leading to subsequent release of calcium from internal stores via PKC signalling [66]. Calcium waves can also be observed within the developing zebrafish brain upon wounding, with these dependent upon glutamate-mediated activation of NMDA receptors. NMDA receptors in turn regulate ATP-dependent recruitment of microglial cells to sites of injury [67]. Traumatic brain injury has long been associated with rapid decreases in extracellular calcium (e.g. Young et al., 1982 [68]; for review see Weber, 2005 [69]) and laser-wounding has been proposed as a useful technique to model this [70]. Indeed, buffering calcium dynamics within damaged neurons in flies can be neuroprotective [71], suggesting this system could uncover novel therapeutic strategies for traumatic brain injury.

In summary, we have shown that increasing the apoptotic cell burden placed on macrophages impairs their normal behaviour, dampening their inflammatory responses to tissue damage. Furthermore, we have uncovered that wound-induced calcium waves spread beyond the epithelium into neighbouring tissues, potentially enabling them to contribute to recruitment of macrophages and subsequent repair processes. How these waves spread and how epithelial and non-epithelial tissues integrate these responses to coordinate inflammation and repair remain key questions for future work. Since specification of macrophages is not perturbed in *repo* mutants, this model will prove useful to examine macrophage-apoptotic cell interactions in vivo in more detail and shed light on cellular interactions that are fundamental for normal development and homeostasis.

## Acknowledgements

We thank Will Wood (University of Edinburgh) for advice and support, Frederico Rodrigues (University of Bristol) for performing preliminary experiments and Karen Plant (University of Sheffield) for technical support. Imaging work was performed in the Wolfson Light Microscopy Facility, using the Perkin Elmer spinning disk (MRC grant G0700091 and Wellcome grant 077544/Z/05/Z) and Nikon A1 confocal/TIRF (Wellcome grant WT093134AIA) microscopes. This work would not be possible without the Bloomington *Drosophila* Stock Centre (NIH P40OD018537) and Flybase (NIH and MRC grants U41 HG000739 and MR/N030117/1, respectively). We thank the *Drosophila* community for sharing *Drosophila* reagents (see Supplementary Table 2). We are grateful to Darren Robinson (Wolfson LMF) and the Fly Facility staff (University of Sheffield) for their assistance and Phil Elks, Simon Johnston, Steve Renshaw and Martin Zeidler (University of Sheffield) for critical reading and feedback on the manuscript. This work was supported by a Sir Henry Dale Fellowship awarded to I.R.E. by Wellcome and The Royal Society (102503/Z/13/Z) and a University of Sheffield Medical School PhD position awarded to I.R.E. and H.G.R.

## Availability of Data and Materials

Fly lines and raw data are available on request from I.R.E.

## References

1. Morioka S, Maueröder C, Ravichandran KS. Living on the Edge: Efferocytosis at the Interface of Homeostasis and Pathology. Immunity. 2019;50: 1149–1162. doi:10.1016/J.IMMUNI.2019.04.018

2. Serhan CN, Savill J. Resolution of inflammation: the beginning programs the end. Nat Immunol. 2005;6: 1191–1197. doi:10.1038/ni1276

3. Bäck M, Yurdagul A, Tabas I, Öörni K, Kovanen PT. Inflammation and its resolution in atherosclerosis: mediators and therapeutic opportunities. Nature Reviews Cardiology. 2019. doi:10.1038/s41569-019-0169-2

4. McCubbrey AL, Curtis JL. Efferocytosis and lung disease. Chest. 2013. pp. 1750–1757. doi:10.1378/chest.12-2413

5. Buchon N, Silverman N, Cherry S. Immunity in Drosophila melanogaster — from microbial recognition to whole-organism physiology. Nat Rev Immunol. 2014;14: 796–810. doi:10.1038/nri3763

6. Banerjee U, Girard JR, Goins LM, Spratford CM. *Drosophila* as a Genetic Model for Hematopoiesis. Genetics. 2019;211: 367–417. doi:10.1534/genetics.118.300223

7. Ratheesh A, Belyaeva V, Siekhaus DE. Drosophila immune cell migration and adhesion during embryonic development and larval immune responses. Current Opinion in Cell Biology. 2015. pp. 71–79. doi:10.1016/j.ceb.2015.07.003

8. Wood W, Jacinto A. Drosophila melanogaster embryonic haemocytes: masters of multitasking. Nat Rev Mol Cell Biol. 2007;8: 542–551. doi:10.1038/nrm2202

9. Evans CJ, Hartenstein V, Banerjee U. Thicker Than Blood. Dev Cell. 2003;5: 673–690. doi:10.1016/S1534-5807(03)00335-6

10. Sears HC, Kennedy CJ, Garrity P a. Macrophage-mediated corpse engulfment is required for normal Drosophila CNS morphogenesis. Development. 2003;130: 3557–3565. doi:10.1242/dev.00586

11. Olofsson B, Page DT. Condensation of the central nervous system in embryonic Drosophila is inhibited by blocking hemocyte migration or neural activity. Dev Biol. 2005;279: 233–243. doi:10.1016/j.ydbio.2004.12.020

12. Defaye A, Evans I, Crozatier M, Wood W, Lemaitre B, Leulier F. Genetic ablation of Drosophila phagocytes reveals their contribution to both development and resistance to bacterial infection. J Innate Immun. 2009;1: 322–334.

13. Stramer B, Wood W, Galko MJ, Redd MJ, Jacinto A, Parkhurst SM, et al. Live imaging of wound inflammation in Drosophila embryos reveals key roles for small GTPases during in vivo cell migration. J Cell Biol. 2005;168: 567–573. doi:10.1083/jcb.200405120

14. Razzell W, Evans IR, Martin P, Wood W. Calcium flashes orchestrate the wound inflammatory response through duox activation and hydrogen peroxide release. Curr Biol. 2013;23: 424–429. doi:10.1016/j.cub.2013.01.058

15. Antunes M, Pereira T, Cordeiro J V., Almeida L, Jacinto A. Coordinated waves of actomyosin flow and apical cell constriction immediately after wounding. J Cell Biol. 2013;202: 365–379. doi:10.1083/jcb.201211039

16. Niethammer P, Grabher C, Look AT, Mitchison TJ. A tissue-scale gradient of hydrogen peroxide mediates rapid wound detection in zebrafish. Nature. 2009;459: 996–999. doi:10.1038/nature08119

17. Yoo SK, Starnes TW, Deng Q, Huttenlocher A. Lyn is a redox sensor that mediates leukocyte wound attraction in vivo. Nature. 2011;480: 109–112. doi:10.1038/nature10632

18. Evans IR, Hu N, Skaer H, Wood W. Interdependence of macrophage migration and ventral nerve cord development in Drosophila embryos. Development. 2010;137: 1625–1633. doi:10.1242/dev.046797

19. Crews ST. *Drosophila* Embryonic CNS Development: Neurogenesis, Gliogenesis, Cell Fate, and Differentiation. Genetics. 2019;213: 1111–1144. doi:10.1534/genetics.119.300974

20. Campbell G, Göring H, Lin T, Spana E, Andersson S, Doe CQ, et al. RK2, a glial-specific homeodomain protein required for embryonic nerve cord condensation and viability in Drosophila. Development. 1994;120: 2957–66. Available: http://www.ncbi.nlm.nih.gov/pubmed/7607085

21. Halter D a, Urban J, Rickert C, Ner SS, Ito K, Travers a a, et al. The homeobox gene repo is required for the differentiation and maintenance of glia function in the embryonic nervous system of Drosophila melanogaster. Development. 1995;121: 317–332.

22. Xiong WC, Okano H, Patel NH, Blendy JA, Montell C. repo encodes a glial-specific homeo domain protein required in the Drosophila nervous system. Genes Dev. 1994. doi:10.1101/gad.8.8.981

23. Shklyar B, Sellman Y, Shklover J, Mishnaevski K, Levy-Adam F, Kurant E. Developmental regulation of glial cell phagocytic function during Drosophila embryogenesis. Dev Biol. 2014;393: 255–269. doi:10.1016/j.ydbio.2014.07.005

24. Bernardoni R, Vivancos V, Giangrande a. Glide/Gcm Is Expressed and Required in the Scavenger Cell Lineage. Dev Biol. 1997;191: 118–130. doi:10.1006/dbio.1997.8702

25. Alfonso TB, Jones BW. gcm2 promotes glial cell differentiation and is required with glial cells missing for macrophage development in Drosophila. Dev Biol. 2002;248: 369–383. doi:10.1006/dbio.2002.0740

26. Trébuchet G, Cattenoz PB, Zsámboki J, Mazaud D, Siekhaus DE, Fanto M, et al. The repo homeodomain transcription factor suppresses hematopoiesis in Drosophila and preserves the glial fate. J Neurosci. 2019. doi:10.1523/JNEUROSCI.1059-18.2018

27. Melcarne C, Lemaitre B, Kurant E. Phagocytosis in Drosophila: From molecules and cellular machinery to physiology. Insect Biochem Mol Biol. 2019;109: 1–12. doi:10.1016/J.IBMB.2019.04.002

28. Kurant E, Axelrod S, Leaman D, Gaul U. Six-Microns-Under Acts Upstream of Draper in the Glial Phagocytosis of Apoptotic Neurons. Cell. 2008;133: 498–509. doi:10.1016/j.cell.2008.02.052

29. Roddie HG, Armitage EL, Coates JA, Johnston SA, Evans IR. Simu-dependent clearance of dying cells regulates macrophage function and inflammation resolution. PLoS Biol. 2019. doi:10.1371/journal.pbio.2006741

30. Weavers H, Evans IR, Martin P, Wood W. Corpse Engulfment Generates a Molecular Memory that Primes the Macrophage Inflammatory Response. Cell. 2016;165. doi:10.1016/j.cell.2016.04.049

31. A-Gonzalez N, Quintana JA, García-Silva S, Mazariegos M, González de la Aleja A, Nicolás-Ávila JA, et al. Phagocytosis imprints heterogeneity in tissue-resident macrophages. J Exp Med. 2017;214: 1281–1296. doi:10.1084/jem.20161375

32. Brückner K, Kockel L, Duchek P, Luque CM, Rørth P, Perrimon N. The PDGF/VEGF receptor controls blood cell survival in Drosophila. Dev Cell. 2004;7: 73–84. doi:10.1016/j.devcel.2004.06.007

33. Stramer B, Moreira S, Millard T, Evans I, Huang CY, Sabet O, et al. Clasp-mediated microtubule bundling regulates persistent motility and contact repulsion in Drosophila macrophages in vivo. J Cell Biol. 2010;189: 681–689.

34. Yoffe KB, Manoukian AS, Wilder EL, Brand AH, Perrimon N. Evidence for engrailed-independent wingless autoregulation in Drosophila. Dev Biol. 1995. doi:10.1006/dbio.1995.1243

35. Ito K, Awano W, Suzuki K, Hiromi Y, Yamamoto D. The Drosophila mushroom body is a quadruple structure of clonal units each of which contains a virtually identical set of neurones and glial cells. Development. 1997.

36. Wodarz A, Hinz U, Engelbert M, Knust E. Expression of crumbs confers apical character on plasma membrane domains of ectodermal epithelia of drosophila. Cell. 1995;82: 67–76. doi:10.1016/0092-8674(95)90053-5

37. Luo L, Joyce Liao Y, Jan LY, Jan YN. Distinct morphogenetic functions of similar small GTPases: Drosophila Drac1 is involved in axonal outgrowth and myoblast fusion. Genes Dev. 1994. doi:10.1101/gad.8.15.1787

38. Lee BP, Jones BW. Transcriptional regulation of the Drosophila glial gene repo. Mech Dev. 2005;122: 849–62. doi:10.1016/j.mod.2005.01.002

39. Chen T-W, Wardill TJ, Sun Y, Pulver SR, Renninger SL, Baohan A, et al. Ultrasensitive fluorescent proteins for imaging neuronal activity. Nature. 2013;499: 295–300. doi:10.1038/nature12354

40. White K, Grether M, Abrams J, Young L, Farrell K, Steller H. Genetic control of programmed cell death in Drosophila. Science (80-). 1994;264: 677–683. doi:10.1126/science.8171319

41. Halfon MS, Gisselbrecht S, Lu J, Estrada B, Keshishian H, Michelson AM. New fluorescent protein reporters for use with thedrosophila gal4 expression system and for vital detection of balancer chromosomes. genesis. 2002;34: 135–138. doi:10.1002/gene.10136

42. Le T, Liang Z, Patel H, Yu MH, Sivasubramaniam G, Slovitt M, et al. A New Family of Drosophila Balancer Chromosomes With a w- dfd-GMR Yellow Fluorescent Protein Marker. Genetics. 2006;174: 2255–2257. doi:10.1534/genetics.106.063461

43. Evans IR, Rodrigues FSLM, Armitage EL, Wood W. Draper/CED-1 Mediates an Ancient Damage Response to Control Inflammatory Blood Cell Migration In Vivo. Curr Biol. 2015;25: 1606–1612. doi:10.1016/j.cub.2015.04.037

44. Schindelin J, Arganda-Carreras I, Frise E, Kaynig V, Longair M, Pietzsch T, et al. Fiji: an open-source platform for biological-image analysis. Nat Methods. 2012;9: 676–682. doi:10.1038/nmeth.2019

45. Sonnenfeld MJ, Jacobs JR. Macrophages and glia participate in the removal of apoptotic neurons from theDrosophila embryonic nervous system. J Comp Neurol. 1995;359: 644–652. doi:10.1002/cne.903590410

46. Freeman MR, Delrow J, Kim J, Johnson E, Doe CQ. Unwrapping glial biology: Gcm target genes regulating glial development, diversification, and function. Neuron. 2003;38: 567–580. doi:10.1016/S0896-6273(03)00289-7

47. Manaka J, Kuraishi T, Shiratsuchi A, Nakai Y, Higashida H, Henson P, et al. Draper-mediated and phosphatidylserine-independent phagocytosis of apoptotic cells by Drosophila hemocytes/macrophages. J Biol Chem. 2004;279: 48466–48476. doi:10.1074/jbc.M408597200

48. Nambu JR, Franks RG, Hu S, Crews ST. The single-minded gene of Drosophila is required for the expression of genes important for the development of CNS midline cells. Cell. 1990;63: 63–75. doi:10.1016/0092-8674(90)90288-P

49. Evans IR, Ghai PA, Urbančič V, Tan KL, Wood W. SCAR/WAVE-mediated processing of engulfed apoptotic corpses is essential for effective macrophage migration in Drosophila. Cell Death Differ. 2013;20: 709–720. doi:10.1038/cdd.2012.166

50. Song Z, McCall K, Steller H. DCP-1, a Drosophila cell death protease essential for development. Science (80-). 1997;275: 536–540. doi:10.1126/science.275.5299.536

51. Moreira S, Stramer B, Evans I, Wood W, Martin P. Prioritization of Competing Damage and Developmental Signals by Migrating Macrophages in the Drosophila Embryo. Curr Biol. 2010;20: 464–470. doi:10.1016/j.cub.2010.01.047

52. Hunter M V., Willoughby PM, Bruce AEE, Fernandez-Gonzalez R. Oxidative Stress Orchestrates Cell Polarity to Promote Embryonic Wound Healing. Dev Cell. 2018;47: 377–387.e4. doi:10.1016/j.devcel.2018.10.013

53. Medina CB, Ravichandran KS. Do not let death do us part: ‘find-me’ signals in communication between dying cells and the phagocytes. Cell Death Differ. 2016;23: 979–989. doi:10.1038/cdd.2016.13

54. Fadok VA, Bratton DL, Konowal A, Freed PW, Westcott JY, Henson PM. Macrophages that have ingested apoptotic cells in vitro inhibit proinflammatory cytokine production through autocrine/paracrine mechanisms involving TGF-beta, PGE2, and PAF. J Clin Invest. 1998;101: 890–898. doi:10.1172/JCI1112

55. Chabaud M, Heuzé ML, Bretou M, Vargas P, Maiuri P, Solanes P, et al. Cell migration and antigen capture are antagonistic processes coupled by myosin II in dendritic cells. Nat Commun. 2015;6: 7526. doi:10.1038/ncomms8526

56. Veltman DM. Drink or drive: competition between macropinocytosis and cell migration. Biochem Soc Trans. 2015;43: 129–132. doi:10.1042/BST20140251

57. Bettman N, Avivi I, Rosenbaum H, Bisharat L, Katz T. Impaired migration capacity in monocytes derived from patients with Gaucher disease. Blood Cells, Mol Dis. 2015. doi:10.1016/j.bcmd.2014.12.003

58. Morell GP, Niaudet P, Jean G, Descamps-Latscha B. Altered oxidative metabolism, motility, and adherence in phagocytic cells from cystinotic children. Pediatr Res. 1985. doi:10.1203/00006450-198512000-00023

59. Aker M, Zimran A, Abrahamov A, Horowitz M, Matzner Y. Abnormal neutrophil chemotaxis in Gaucher disease. Br J Haematol. 1993. doi:10.1111/j.1365-2141.1993.tb08270.x

60. Berg RD, Levitte S, O’Sullivan MP, O’Leary SM, Cambier CJ, Cameron J, et al. Lysosomal Disorders Drive Susceptibility to Tuberculosis by Compromising Macrophage Migration. Cell. 2016;165. doi:10.1016/j.cell.2016.02.034

61. Edwards-Jorquera SS, Bosveld F, Bellaïche YA, Lennon-Duménil A-M, Glavic Á. Trpml controls actomyosin contractility and couples migration to phagocytosis in fly macrophages. J Cell Biol. 2020;219. doi:10.1083/jcb.201905228

62. Kato K, Losada-Perez M, Hidalgo A. Gene network underlying the glial regenerative response to central nervous system injury. Dev Dyn. 2018;247: 85–93. doi:10.1002/dvdy.24565

63. Restrepo S, Basler K. Drosophila wing imaginal discs respond to mechanical injury via slow InsP3R-mediated intercellular calcium waves. Nat Commun. 2016;7: 12450. doi:10.1038/ncomms12450

64. Clark AG, Miller AL, Vaughan E, Yu H-YE, Penkert R, Bement WM. Integration of single and multicellular wound responses. Curr Biol. 2009;19: 1389–95. doi:10.1016/j.cub.2009.06.044

65. Xu S, Chisholm AD. A Gαq-Ca^2+^ signaling pathway promotes actin-mediated epidermal wound closure in C. elegans. Curr Biol. 2011;21: 1960–7. doi:10.1016/j.cub.2011.10.050

66. de Oliveira S, López-Muñoz A, Candel S, Pelegrín P, Calado Â, Mulero V. ATP modulates acute inflammation in vivo through dual oxidase 1-derived H2O2 production and NF-κB activation. J Immunol. 2014;192: 5710–9. doi:10.4049/jimmunol.1302902

67. Sieger D, Moritz C, Ziegenhals T, Prykhozhij S, Peri F. Long-Range Ca2+ Waves Transmit Brain-Damage Signals to Microglia. Dev Cell. 2012;22: 1138–1148. doi:10.1016/j.devcel.2012.04.012

68. Young W, Yen V, Blight A. Extracellular calcium ionic activity in experimental spinal cord contusion. Brain Res. 1982. doi:10.1016/0006-8993(82)90677-1

69. Weber J. Calcium Homeostasis Following Traumatic Neuronal Injury. Curr Neurovasc Res. 2005. doi:10.2174/1567202043480134

70. Shannon EK, Stevens A, Edrington W, Zhao Y, Jayasinghe AK, Page-McCaw A, et al. Multiple Mechanisms Drive Calcium Signal Dynamics around Laser-Induced Epithelial Wounds. Biophys J. 2017;113: 1623–1635. doi:10.1016/j.bpj.2017.07.022

71. Avery MA, Rooney TM, Pandya JD, Wishart TM, Gillingwater TH, Geddes JW, et al. WldS prevents axon degeneration through increased mitochondrial flux and enhanced mitochondrial Ca2+ buffering. Curr Biol. 2012;22: 596–600. doi:10.1016/j.cub.2012.02.043

