## Supplementary Figure, Table and Movie Legends for "Overexposure to apoptosis via disrupted glial specification perturbs *Drosophila* macrophage function and reveals roles of the CNS during injury"

### Supplementary Figure 1. Maintenance of elevated calcium levels at wound sites

(a-b) *w^1118^;;da-GAL4,UAS-GCaMP6M* embryos 1 hour (a) and 2 hours (b) post-wounding showing elevated calcium levels at wound sites. Scale bars represent 100μm.

### Supplementary Table 1. University of Sheffield fly food ingredients and suppliers

### Supplementary Table 2. Genotypes and sources of alleles used in this study

### Supplementary Movie 1. Defective inflammatory responses to injury in the absence of *repo* function

Response of GFP and red stinger-labelled macrophages to injury in control and *repo* mutant embryos. Embryos wounded on the ventral midline at stage 15. Scale bar represents 20μm. Genotypes are *w^1118^;srp-GAL4,UAS-red stinger/srp-GAL4,UAS-GFP* (control) and *w^1118^;srp-GAL4,UAS-red stinger/srp-GAL4,UAS-GFP;P{PZ}repo^03702^* (*repo*).

### Supplementary Movie 2. Epithelial calcium responses to injury in stage 15 control and *repo* mutant embryos

Movies of epithelial cell calcium responses to injury at stage 15 in control and *repo* mutant embryos. Calcium imaged via *w^1118^;;da-GAL4,UAS-GCaMP6M* and projections assembled from superficial slices of z-stacks. Embryos wounded on the ventral midline. Scale bars show 20μm. Movies correspond to stills shown in Figure 7a-b.

### Supplementary Movie 3. Calcium responses of non-epithelial tissue following wounding

Movie of calcium responses to injury at stage 15 at different depths within the embryo. Calcium dynamics visualised via ubiquitous expression of GCaMP6M (*w^1118^;;da-GAL4,UAS-GCaMP6M*) following wounding on the ventral midline. Movie assembled from slices corresponding to superficial z-slices of the image stack (epithelial cells, 0-5μm from the surface of the embryo; top panel), slices of the ventral half of the VNC (10-20μm from the surface of the embryo; central panel) and the dorsal half of the VNC (20-30μm from the surface of the embryo; bottom panel). Scale bar shows 20μm. Movie corresponds to stills shown in Figure 7e-g.

### Supplementary Movie 4. Neuronal calcium responses to injury

Movie of neuronal calcium response to injury at stage 15. Calcium imaged in neurons via *w^1118^;;elav-GAL4,UAS-GCaMP6M*. Embryo wounded laterally with respect to the ventral midline. Scale bar shows 20μm. Movie corresponds to stills shown in Figure 8d.

### Supplementary Movie 5. Calcium responses of glial cells to injury in control and *repo* mutant embryos

Movies of glial calcium responses to injury at stage 15 in control and *repo* mutant embryos. Calcium imaged via *repo-GAL4,UAS-GCaMP6M*. Embryos wounded on the ventral midline. Scale bars show 20μm. Movies correspond to stills shown in Figure 8e-f. Genotypes are *w^1118^;repo-GAL4,UAS-GCaMP6M* (control) and *w^1118^;repo-GAL4,UAS-GCaMP6M;P{PZ}repo^03702^* (*repo*).
