## Supplementary figures and images for "Overexposure to apoptosis via disrupted glial specification perturbs *Drosophila* macrophage function and reveals roles of the CNS during injury"

### Supplementary Figure 1

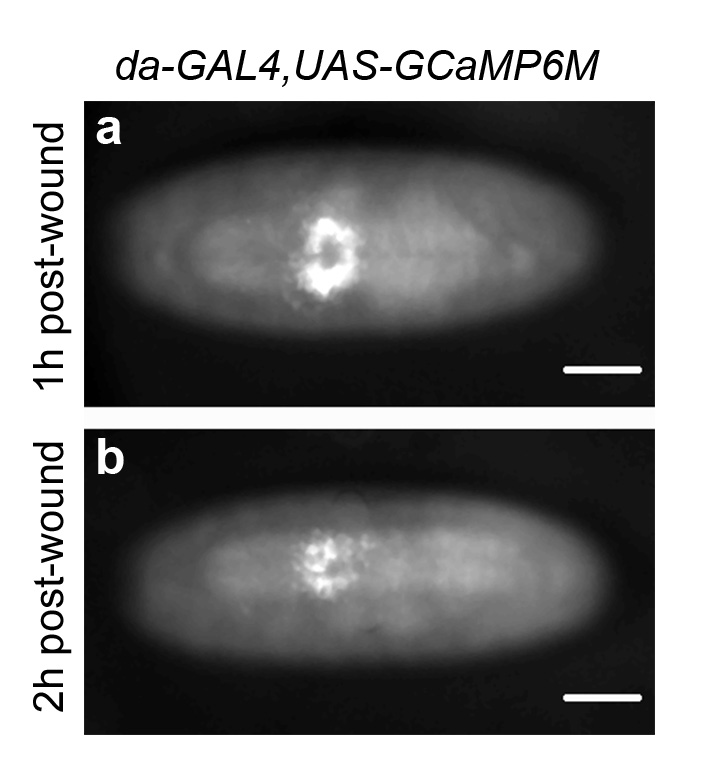
