## Supplementary Table 2 for "Overexposure to apoptosis via disrupted glial specification perturbs *Drosophila* macrophage function and reveals roles of the CNS during injury"

| Figure panel(s) | Label | Genotype | Source | Location |
|---|---|---|---|---|
| **Figure 1a-b** | control | *w[1118];srp-GAL4,UAS-GFP/+;crq-GAL4,UAS-GFP/+* | *srp-GAL4,UAS-GFP* and *crq-GAL4,UAS-GFP* from Will Wood | University of Edinburgh, UK |
| **Figure 1c-d** | control | *w[1118];;crq-GAL4,UAS-GFP* | | |
| **Figure 1e** | *repo*$^{03702}$ | *w[1118];;P{PZ}repo[03702],crq-GAL4,UAS-GFP* | recombinant of *P{PZ}repo[03702]*, a Bloomington stock, n.b. *ry[506]* recombined off | Bloomington Drosophila Stock Centre, USA |
| **Figure 2a** | control | *w[1118];srp-GAL4,UAS-GFP/srp-GAL4,UAS-red stinger* | *UAS-red stinger* from Brian Stramer | KCL, UK |
| **Figure 2b** | *repo*$^{03702}$ | *w[1118];srp-GAL4,UAS-GFP/srp-GAL4,UAS-red stinger;ry[506],P{PZ}repo[03702]* | | |
| **Figure 2c,e** | control | *w[1118];;crq-GAL4,UAS-GFP* | | |
| **Figure 2d-e** | *repo*$^{03702}$ | *w[1118];;P{PZ}repo[03702],crq-GAL4,UAS-GFP* | | |
| **Figure 3a-c,e** | control | *w[1118];;crq-GAL4,UAS-GFP* | | |
| **Figure 3a-b,d-e** | *repo*$^{03702}$ | *w[1118];;P{PZ}repo[03702],crq-GAL4,UAS-GFP* | | |
| **Figure 3a-b** | *simu*$^{2}$ | *w[1118];simu[2];crq-GAL4,UAS-GFP* | *simu[2]* from Estee Kurant | University of Haifa, Israel |
| **Figure 3a-b** | *simu;repo* | *w[1118];simu[2];P{PZ}repo[03702],crq-GAL4,UAS-GFP* | | |
| **Figure 4a,e** | control | *w[1118];srp-GAL4,UAS-GFP/srp-GAL4,UAS-red stinger* | | |
| **Figure 4b,e** | *Df(3L)H99* | *w[1118];srp-GAL4,UAS-GFP/srp-GAL4,UAS-red stinger;Df(3L)H99* | | |
| **Figure 4c,e** | *repo*$^{03702}$ | *w[1118];srp-GAL4,UAS-GFP/srp-GAL4,UAS-red stinger;P{PZ}repo[03702]* | | |
| **Figure 4d-e** | *Df(3L)H99,repo*$^{03702}$ | *w[1118];srp-GAL4,UAS-GFP/srp-GAL4,UAS-red stinger;Df(3L)H99,P{PZ}repo[03702* | recombinant of *Df(3L)H99*, a Bloomington stock | Bloomington Drosophila Stock Centre, USA |
| **Figure 5a,c-f** | control | *w[1118];;crq-GAL4,UAS-GFP* | | |
| **Figure 5b-f** | *repo*$^{03702}$ | *w[1118];;P{PZ}repo[03702],crq-GAL4,UAS-GFP* | | |
| **Figure 6a,e-f** | control | *w[1118];srp-GAL4,UAS-GFP/srp-GAL4,UAS-red stinger* | | |
| **Figure 6b,e-f** | *Df(3L)H99* | *w[1118];srp-GAL4,UAS-GFP/srp-GAL4,UAS-red stinger;Df(3L)H99* | | |
| **Figure 6c,e-f** | *repo*$^{03702}$ | *w[1118];srp-GAL4,UAS-GFP/srp-GAL4,UAS-red stingerP{PZ};repo[03702]* | | |
| **Figure 6d-f** | *Df(3L)H99,repo*$^{03702}$ | *w[1118];srp-GAL4,UAS-GFP/srp-GAL4,UAS-red stinger;Df(3L)H99,P{PZ}repo[03702]* | | |
| **Figure 7a,c-g** | control | *w[1118];;da-GAL4,UAS-GCaMP6M* | recombinant of *UAS-GCaMP6M* (a Bloomington stock) and *da-GAL4* from Alex Whitworth | Bloomington Drosophila Stock Centre, USA/University of Cambridge, UK |
| **Figure 7b-d** | *repo*$^{03702}$ | *w[1118];;P{PZ}repo[03702],da-GAL4,UAS-GCaMP6M* | | |
| **Figure 8a,c** | control | *w[1118];e22c-GAL4,UAS-GCaMP6M* | recombinant of *e22c-GAL4*, a Bloomington stock | Bloomington Drosophila Stock Centre, USA |
| **Figure 8b-c** | *repo*$^{03702}$ | *w[1118];e22c-GAL4,UAS-GCaMP6M;P{PZ}repo[03702]* | | |
| **Figure 8d** | control | *w[1118],elav-GAL4/w[1118];UAS-GCaMP6M/+* | *w[1118],elav-GAL4*, a Bloomington stock | Bloomington Drosophila Stock Centre, USA |
| **Figure 8e,g-i** | control | *w[1118];repo-GAL4,UAS-GCaMP6M* | recombinant of *repo-GAL4* from Estee Kurant | University of Haifa, Israel |
| **Figure 8f,h-i** | *repo*$^{03702}$ | *w[1118];repo-GAL4,UAS-GCaMP6M;P{PZ}repo[03702]* | | |
| **Supplementary movie 1** | control | *w[1118];srp-GAL4,UAS-GFP/srp-GAL4,UAS-red stinger* | | |
| **Supplementary movie 1** | *repo*$^{03702}$ | *w[1118];srp-GAL4,UAS-GFP/srp-GAL4,UAS-red stinger;P{PZ}repo[03702]* | | |
| **Supplementary movie 2** | control | *w[1118];;da-GAL4,UAS-GCaMP6M* | | |
| **Supplementary movie 2** | *repo*$^{03702}$ | *w[1118];;P{PZ}repo[03702],da-GAL4,UAS-GCaMP6M* | | |
| **Supplementary movie 3** | control | *w[1118];;da-GAL4,UAS-GCaMP6M* | | |
| **Supplementary movie 4** | control | *w[1118],elav-GAL4/w[1118];UAS-GCaMP6M/+* | | |
| **Supplementary movie 5** | control | *w[1118];repo-GAL4,UAS-GCaMP6M* | | |
| **Supplementary movie 5** | *repo*$^{03702}$ | *w[1118];repo-GAL4,UAS-GCaMP6M;P{PZ}repo[03702]* | | |
| **Supplementary figure 1a-b** | control | *w[1118];;da-GAL4,UAS-GCaMP6M* | | |
